# Hebbian activity only temporarily stabilizes synaptic transmission at CA3-CA1 synapses in the developing hippocampus

**DOI:** 10.1101/2024.01.10.574846

**Authors:** Joakim Strandberg, Bengt Gustafsson

## Abstract

Prolonged low frequency (0.05-1 Hz) stimulation of previously non-stimulated (naive) CA3-CA1 synapses in the developing hippocampus results in a profound synaptic depression explained by a postsynaptic AMPA silencing. It has been proposed that Hebbian activity can stabilize the synapses by preventing such depression. Using field recordings, we have examined to which extent strong repeated high frequency tetanization simulating Hebbian activity results in such prevention. The tetanization resulted within minutes in a field EPSP potentiation to 150-170% of the naive field EPSP level which remained unaltered if stimulation was suspended. If test pulse stimulation (0.2 Hz) was allowed to continue after the tetanization the field EPSP continuously decreased and was after 2700 stimuli depressed by 75% from the potentiated level. This depression did not differ in relative terms from that induced in naive synapses (by 82% from the naive level). The long-lasting component of this depression revealed by a subsequent 30 min stimulus interruption (by 59% from the potentiated level) did not differ from that of naive synapses (by 66% from the naive level). This equal relative degree of depression of tetanized and naive synapses was also observed following 2700 stimuli at 1 Hz. On the other hand, when examined at earlier time points during the test pulse stimulation (e.g. after 400-900 stimuli) tetanized synapses were less depressed than naive synapses, and the long-lasting depression after 900 stimuli at 1 Hz was only half that observed in naive synapses. This effect of tetanization was observed independently of whether the 1 Hz stimulation was commenced 15 min or 2 hours after the tetanization. In conclusion, while a strong preceding tetanization results in a partial stabilization of transmission at CA3-CA1 synapses in the developing hippocampus, this effect appears only temporary. This temporary effect is not linked to time after tetanization but to the number of low frequency stimuli given.

## INTRODUCTION

During brain development synapses are continuously generated and thereafter selected in an activity dependent manner to establish an appropriate mature pattern of synaptic connectivity. While little is known about the mechanisms that initiate the break-up of established synapses or that protect synapses from elimination, activity dependent synaptic plasticity which may differ in important respects from that in the mature nervous system are likely of critical importance. In fact, when CA3-CA1 synapses are subjected to low frequency stimulation in the developing, but not in the mature, hippocampus, they become considerably depressed even when stimulated only a few times per minute. This depression was found to be initiated to much the same extent per stimulus over a large frequency range (0.05-1 Hz) and to be explained by AMPA silencing (Abrahamsson et al., 2007; Strandberg et al., 2009; Xiao et al., 2004). In the developing hippocampus even rather sparse activity may thus result in many CA3-CA1 synapses losing their AMPA receptors (AMPARs), an absence that if occurring too frequently may be an initial step towards synapse elimination (Bastrikova et al., 2008; Becker et al., 2008; Kamikubo et al., 2006).

The depression induced by low frequency stimulation can reverse in 20-30 min by a stimulus interruption or reverse by high frequency tetanization (Hebbian activity) (Abrahamsson et al., 2007, 2008; Xiao et al., 2004). When the depression is reversed by stimulus interruption a resumed low frequency stimulation easily depresses the synapses again (Abrahamsson et al., 2008). That is, while presynaptic silence can allow a synapse to maintain its AMPARs, these receptors can easily be lost again if the synapse is activated, even sporadically. In contrast, when the depression is reversed after a high-frequency tetanization the synapses appear to stabilize at the naive level present prior to the initiation of stimulation of these synapses, a level when essentially no synapses are AMPA silent (Abrahamsson et al., 2008; Xiao et al., 2004). Hebbian activity in the 2^nd^ postnatal week thus seems to produce plasticity that differs distinctly from that in the more mature brain in that no actual lasting potentiation takes place. Instead, Hebbian activity de-depresses the synapses and appears to transform the synapses into a state in which they are stable towards the depressive effect of low frequency stimulation (Groc et al., 2006). Such a transformation would protect the synapses from easily losing their AMPARs and possibly be protective against later elimination.

Synaptic plasticity in the developing brain may thus have the character that synapses are easily silenced and possibly later eliminated if they are active without participating in Hebbian activity but are on the other hand protected against silencing after having participated in Hebbian activity. However, whether such a protection is permanent or only temporary is unknown since the stabilization of the naive AMPA signaling level after tetanization has only been followed up to 60 min post-tetanus. In the present study we have used more prolonged low frequency stimulation following high frequency tetanization to examine in what manner participation in Hebbian activity alters the sensitivity of CA3-CA1 synapses in the developing hippocampus to low frequency stimulation.

## METHODS

Most experimental details have previously been described in (Strandberg et al., 2009) and (Strandberg & Gustafsson, 2011). In brief, experiments were performed on hippocampal slices from 8–12-day-old Wistar rats kept and killed in accordance with the guidelines of the Gothenburg ethical committee for animal research. The brain was removed, placed in an ice–cold solution and transverse hippocampal slices (400 µm thick) were cut with a vibratome. After typically 2-5 hours of storage at 25ºC a single slice was transferred to a recording chamber where it was kept submerged in a constant flow (∼2 ml per minute) at ∼30ºC. The perfusion ACSF contained (in mM): 124 NaCl, 3 KCl, 4 CaCl_2_, 4 MgCl_2_, 26 NaHCO_3_, 1.25 NaH_2_PO_4_, and 10 *D*–glucose. Picrotoxin (100 µM) was always present in the perfusion ACSF to block GABA_A_ receptor-mediated activity.

Electrical stimulation of Schaffer collateral afferents was carried out in the stratum radiatum delivered through a tungsten microelectrode (resistance ∼0.1 MΩ) insulated except at its tip. Usually, two stimulating electrodes were positioned on either side of the recording electrode to provide for two independent synaptic inputs to the same dendritic region. Field EPSP recordings were made by means of a glass micropipette (∼2 MΩ, filled with 1 M NaCl) in the stratum radiatum. The field EPSP magnitude was measured, using linear regression over the first 0.8 ms, as the initial slope of the field EPSP rising phase. With the stimulation intensities used (20-50 µA) the naive field EPSPs were generally subthreshold for spike generation. The magnitude of the presynaptic volley was estimated by linear regression of the negative slope of the initial positive-negative deflection. The field EPSP slope measurements in each experiment were linearly adjusted for by changes in the magnitude of the presynaptic volley. For validation of this “volley correction” of the field EPSP, see (Strandberg et al., 2009). For the estimation of the naive level of synaptic transmission only the very 1^st^ evoked field EPSP was used while for the estimation of the amount of depression attained at a certain time during the stimulation the average of 20 field EPSPS was used. Because the field EPSP obtained after stimulus interruption could be quite small, the field EPSP level after stimulus interruption was calculated as the average of the first three field EPSPs obtained when stimulation was resumed.

The high-frequency tetanization protocol consisted of three tetanization events 7.5 min apart, each event consisting of three 20-impulse 50 Hz tetani at 0.1 Hz. To ascertain that the induction conditions were strong enough to fulfill the cooperativity (Hebbian) requirement for LTP induction which may otherwise constitute a problem in the developing hippocampus (Harris & Teyler, 1984; Liao & Malinow, 1996) the tetanization of the synaptic test input was always made in conjunction with tetanization of a second conditioning synaptic input (as in (Abrahamsson et al., 2008)). Data are expressed as means ± SEM. Statistical significance for independent samples was evaluated using Student’s t–test.

### Drugs

Chemicals were from Sigma–Aldrich (Stockholm, Sweden).

## RESULTS

### Does high frequency tetanization stabilize synaptic transmission at the naive level?

High-frequency tetanization of previously non-stimulated (naive) CA3-CA1 synapses in slices from the 2^nd^ postnatal week rat hippocampus does not result in any stable potentiation of the field EPSP. Instead, when followed by test pulse stimuli (0.05-0.2 Hz) for 30-60 min post-tetanus there is only a transient (5-15 min) enhancement above the naive level before the field EPSP seemingly stabilizes at the naive level (Abrahamsson et al., 2008). In the present experiments we have used this procedure of applying tetanization at the naive level and have thereafter exposed the synapses to very prolonged test pulse (0.2 Hz) stimulation (2700 stimuli). Figure 1A shows that after applying our strong tetanization protocol (see Methods and text-Fig. 1) the field EPSP was after the 3^rd^ tetanization event potentiated to 151% (± 7.3%, n = 6) of the naive level and thereafter decayed back towards the naive level. However, as shown in Figure 1B, the field EPSP did not stabilize at the naive level but decreased continuously during the ∼ 4-hour recording period to 37% of the naive level, i.e., to a depression of 63 ± 2.6% from the naive level (n = 6). Thus, not even strong Hebbian activity can stabilize the naive AMPA signalling level at CA3-CA1 synapses in the developing hippocampus.

**Figure 1.**
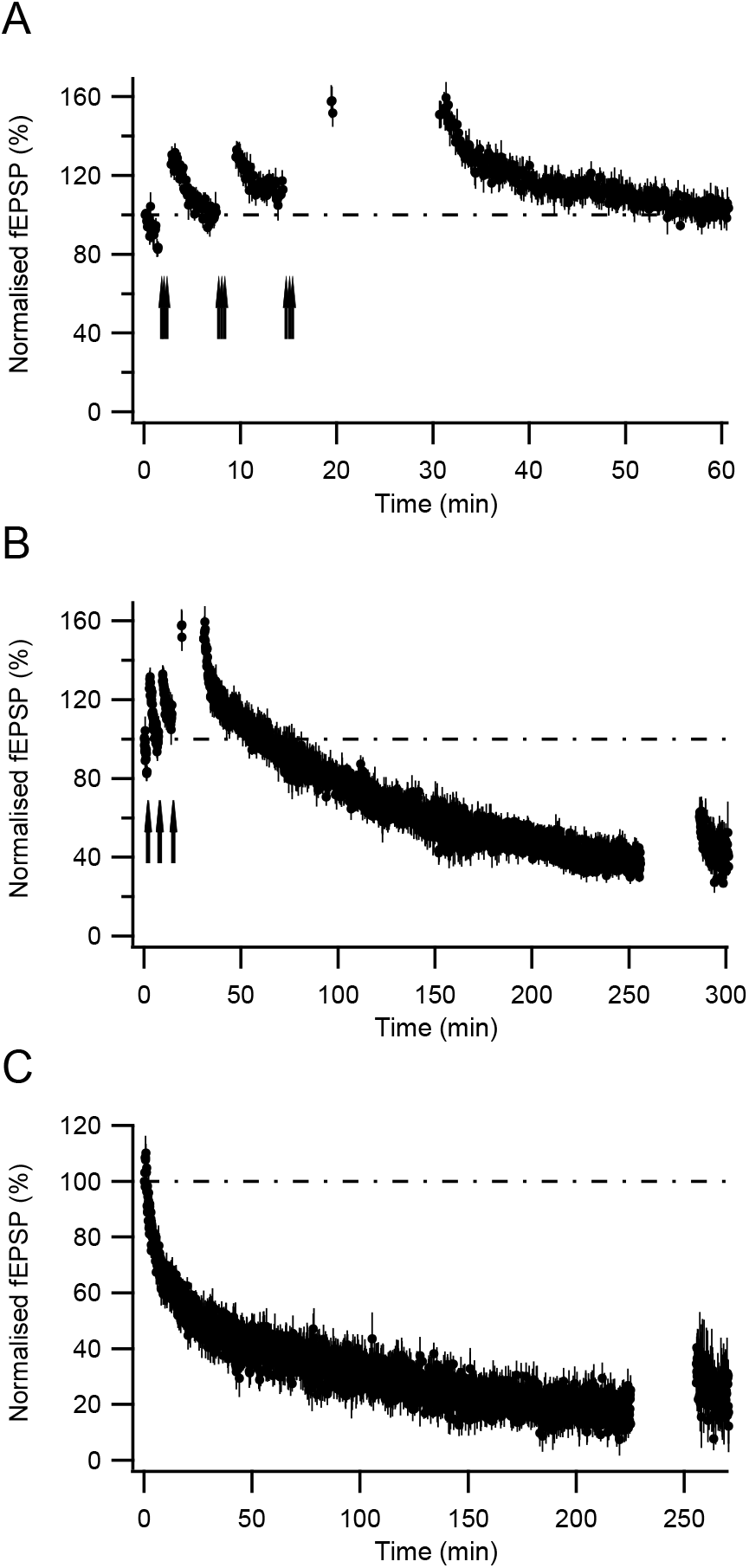
Effect of high frequency tetanization on test pulse-induced depression. A, after 20 stimuli at 0.2 Hz to previously non-stimulated (naive) synapses to establish the naive field EPSP level, three 20-impulse 50 Hz tetani were given at 0.05 Hz (thick arrow) after which the 0.2 Hz stimulation was resumed. This tetanization event was repeated twice with 7.5 min interval. After the third tetanization event the test pulse stimulation (0.2 Hz) was first given three times 5 min after the tetanization and thereafter resumed 15 min after the tetanization. Note that the field EPSP did not decay until the stimulation was resumed 15 min after tetanization (n = 6 experiments). B, same as in A, but on a longer time scale. Note that the field EPSP decreases substantially below the naive level (100%). After 2700 stimuli, test pulse stimulation was interrupted for 30 min to allow the depression to reverse after which the test pulse stimulation was resumed. C, test pulse stimulation (0.2 Hz) applied to previously naive synapses (n = 5 experiments).

Test pulse stimulation of naive CA3-CA1 synapses results in both long-lasting depression and in depression that reverses within a subsequent 30 min period of stimulus interruption (Strandberg & Gustafsson, 2011). The 0.2 Hz-induced depression of the tetanized synapses described above also partially reversed leaving a long-lasting depression of 38 ± 5.8% from the naive level (n = 6) (Fig. 1B). For comparison, when we exposed naive synapses to a similar prolonged 0.2 Hz stimulation (2700 stimuli) this stimulation resulted in a depression of 82 ± 1.8% from the naive level (n = 5) and in a long-lasting depression of 66 ± 4.6% from the naive level (Fig. 1C). Tetanized synapses were thus less depressed than the naive synapses both with respect to the final level of depression reached during the stimulation (63% vs 82%, p < 0.001) and to the long-lasting depression (38% vs 66%, p < 0.02), indicating a partial stabilizing action of Hebbian activity (Fig. 2A).

**Figure 2.**
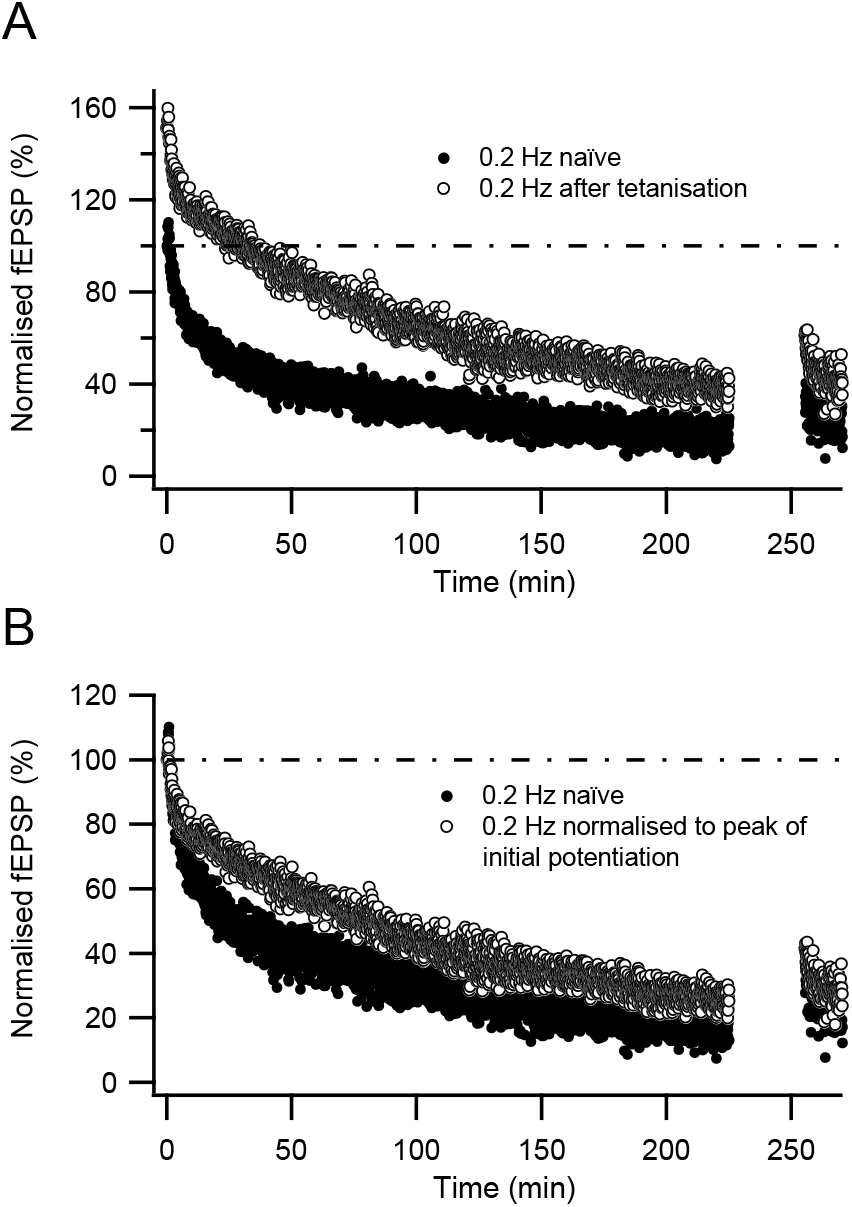
Comparison between test pulse-induced depression of naive and tetanized synapses. A, the depressions shown in figure 1B and C, respectively, are plotted superimposed using the naive level as reference level (100%) for both curves. B, same plot as in A, but the depression of the tetanized synapses is plotted using the post-tetanus-naive level as reference level (100%).

### Which is the “naive” level after the tetanization?

The above comparison between the 0.2 Hz-induced depressions obtained before and after tetanization assumes that the initial potentiation is a transient phenomenon whose impact on the field EPSP is essentially over within 15 min of stimulation and which does not reverse during the stimulus interruption. Figure 1A shows that the prolonged 0.2 Hz stimulation was not initiated immediately after the 3^rd^ tetanization event but that 15 min of stimulus interruption were given allowing the initial potentiation to largely decline (note the transient character of the initial potentiation following the 1^st^ tetanization event). As indicated in figure 1A, however, the field EPSP was as potentiated when the stimulation was resumed after the 15 min of stimulus interruption as when measured following three 0.2 Hz stimuli delivered ∼ 3 min after the 3^rd^ tetanization event. In other experiments (see below) we used a 2-hour stimulus interruption after the tetanization without observing any decrease in the field EPSP. The tetanization-induced potentiation above the naive level is thus not transient by itself but declines in a stimulation dependent manner with a time course that becomes more prolonged with the successive tetanization events (Fig. 1A). To assess the stabilizing action of Hebbian activity on stimulus-induced depression we have therefore also normalized the depression observed after tetanization to the very first field EPSP magnitude obtained when stimulation was resumed 15 min after the 3^rd^ tetanization event. We will hereafter refer to this field EPSP magnitude as the post-tetanus-naive level. Using this normalization procedure, the 0.2 Hz stimulation of tetanized synapses resulted in a depression and in a long-lasting depression of 75 ± 2.7% and 58 ± 5.8% from the post-tetanus-naive level, respectively (Fig. 2B). As evaluated from this post-tetanus-naive level the tetanized synapses thus are as depressed percent wise as the naive synapses both with respect to the depression reached during the stimulation (75% vs 82%, p = 0.06) and to the level of long-lasting depression (58% vs 66%, p > 0.40). Moreover, the reversible component of the depression now becomes proportionally similar at tetanized synapses (17 ± 3.3%) as at the naive synapses (18 ± 1.7%). However, when measured at earlier time points during the stimulation, such as ∼ 30 min after tetanization the tetanized synapses were significantly less depressed (34 ± 3.7% vs 54 ± 2.9%, p < 0.005). Thus, even when evaluated using this normalization it appears that tetanization results in a partial stabilization of the AMPA signalling, albeit more temporarily. In fact, while it takes ∼ 30 min to obtain a depression of 54% from the naive level at naive synapses it takes about three times as long to obtain the same amount of depression from the post-tetanus-naive level at the tetanized synapses (Fig. 2B).

### Does high frequency tetanization affect 1 Hz-induced depression?

We next determined to what extent Hebbian activity also might exert some stabilizing action on the depression induced by the common LTD-inducing 1 Hz stimulation. When applied to naive synapses 900 stimuli at 1 Hz fail to produce significantly more depression than the same number of stimuli at 0.05-0.2 Hz (Strandberg et al., 2009). To determine whether this is true also for tetanized synapses we first examined to what extent this equal potency of 1 Hz and test pulse stimulation to depress naive synapses also extends to the depression induced by 2700 stimuli. We therefore applied 2700 stimuli at 1 Hz to naive synapses which resulted in a depression and a long-lasting depression of 89 ± 3.0% (n = 5) and of 51% ± 4.6% (n = 5) from the naive level, respectively (Fig. 3A). While the 1 Hz stimulation compared to the 0.2 Hz stimulation thus on average produced a somewhat greater depression (89% vs 82%) and a somewhat smaller long-lasting depression (51% vs 66%) none of these differences were statistically significant. However, the reversal from the 1 Hz-induced depression (38 ± 3.1% of the naive level) was significantly larger (p < 0.004) than the reversal from the 0.2 Hz-induced depression (18 ± 1.7% of the naive level). In fact, while prolongation of 1 Hz stimulation of naive synapses from 900 to 2700 stimuli resulted in an increase of the depression from 60% (Strandberg & Gustafsson, 2011) to 89% from the naive level it did not lead to any significant increase in long-lasting depression (45% vs 51%, p = 0.19). In contrast, a similar prolongation of the 0.2 Hz stimulation of naive synapses resulted in a greater depression (63% vs 82%) but also to an even greater increase of the long-lasting depression (38% vs 66%). Thus, while the equal potency of 1 Hz and 0.2 Hz stimulation to depress naive synapses largely holds true also with respect to this more prolonged stimulation, the 1 Hz stimulation is considerably less effective than the 0.2 Hz stimulation to convert the additional depression into a long-lasting depression.

**Figure 3.**
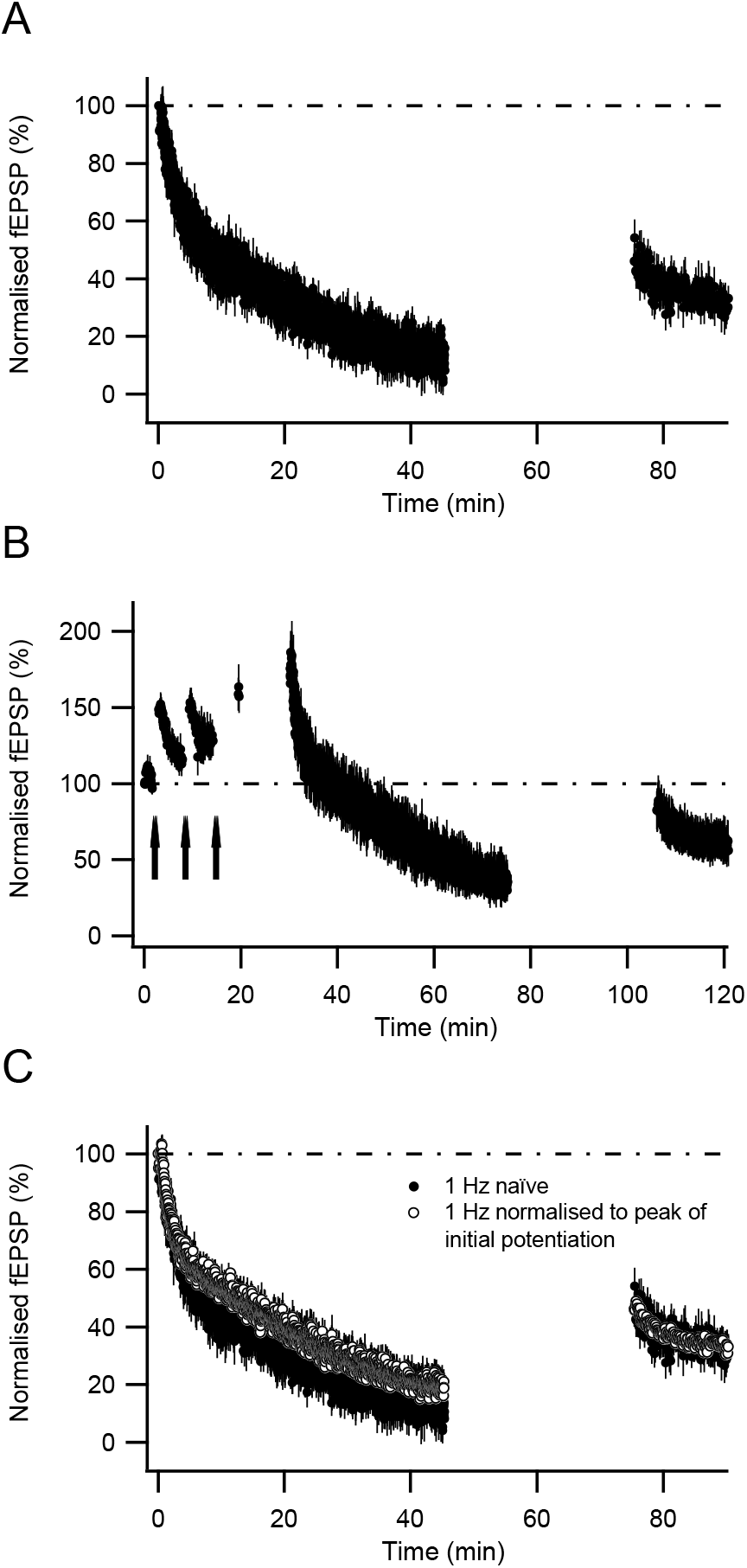
Effect of high frequency tetanization on 1 Hz-induced depression. A, 2700 stimuli at 1Hz was applied to previously naive synapses. After 30 min of stimulus interruption to allow the depression to reverse, 0.2 Hz test pulse stimulation was given (n = 5 experiments). B, same procedure as in Fig. 1 A, B but the stimulation started 15 min after the third tetanization event was at 1 Hz (n = 5 experiments). C, the depressions shown in A and B are plotted superimposed using the post-tetanus-naive level as reference level (100%) for the depression of the tetanized synapses.

We thereafter applied the 2700 stimuli at 1 Hz to tetanized synapses which resulted in a depression and a long-lasting depression of 65 ± 9.8% and of 17 ± 14% (n = 5) from the naive level, respectively, and in a reversible depression of 47 ± 5.5% of the naive level (Fig. 3B). Thus, 1 Hz stimulation failed to produce more depression than the 0.2 Hz stimulation also when applied to tetanized synapses. Moreover, like the naive synapses the 1Hz stimulation of tetanized synapses resulted in a significantly greater reversible depression than the 0.2 Hz stimulation (25 ± 3.9% of the naive level, p < 0.01). As evaluated from the post-tetanus-naive level, the 1 Hz stimulation resulted in a depression and a long-lasting depression of 81 ± 4.0% and of 54 ± 4.4% from that level, respectively, and a reversible depression of 27 ± 2.1% of that level. Thus, also when evaluated in this manner the 1 Hz stimulation failed to produce significantly more depression than the 0.2 Hz stimulation and the reversal of depression was significantly greater (p < 0.04).

To evaluate the possible stabilizing action of Hebbian activity against 1 Hz-induced depression we compared the depression induced at naïve synapses from the naive level with that induced at tetanized synapses from the post-tetanus-naive level (Fig. 3C). Like the 0.2 Hz-induced depression neither the depression (81%) nor the long-lasting depression (54%) differed significantly from the corresponding depressions of the naive synapses (89% and 51%, respectively). On the other hand, the reversal from depression was significantly smaller for the tetanized synapses than for the naive synapses (27% vs 38%, p < 0.03) indicating that the relative failure of the prolonged 1Hz stimulation to result in long-lasting rather than in reversible depression is less pronounced after tetanization. When measured after ∼ 6 min of stimulation at 1 Hz, i.e., after a similar number of stimuli as the early measurement using 0.2 Hz stimulation, the tetanized synapses were less depressed than the naive synapses (40 ± 3.3% vs 48 ± 2.6%) but this difference did not reach statistical significance (p = 0.08).

### Does the 1 Hz-induced long-lasting depression of tetanized synapses depend on time after tetanization?

Since tetanization failed to induce any clear stabilizing effect on the amount of depression produced during on-going 1 Hz stimulation we next examined whether it may have affected the amount of long-lasting depression induced by this stimulation. We have previously shown that 900 stimuli at 1 Hz of naive synapses results in a depression of 62% from the naive level, subdivided in reversible and long-lasting depression of 17% and 45% from the naive level, respectively (Strandberg & Gustafsson, 2011). In the present experiments we applied this stimulation 15 min or 2 hours post-tetanus, respectively, in order both to compare the resulting depressions against each other and to compare them to the depression induced at naive synapses. The two hours of stimulus interruption did not affect the field EPSP since when stimulation was resumed after these two hours its magnitude (167 ± 6.4%, n = 5) was no different from that observed when examined by a few 0.2 Hz stimuli 4 min after the 3rd tetanization (169 ± 4.7%, n = 5, p = 0.77). Figure 4 shows that when normalized to the post-tetanus-naive level the depressions induced by 900 stimuli at 1 Hz applied 15 min and 2 hours post-tetanus, respectively, were quite super imposable and did not differ significantly with respect to either depression, long-lasting depression, or reversible depression. Compared to the naive synapses the depression after 900 stimuli was however somewhat smaller (50 ± 3.3%, n = 10, p < 0.002) (combined data of both 15 min and 2 hours post-tetanus). In contrast, the long-lasting depression was substantially smaller (24 ± 5.3%, n = 8, p < 0.005) and the reversible depression substantially larger (28 ± 3.2%, n = 8, p < 0.02) compared to the naive synapses. Thus, tetanization provides for a temporary stabilization also against 1 Hz stimulation, this effect however being most evident with respect to the ability of the stimulation to convert depression into a long-lasting depression.

**Figure 4.**
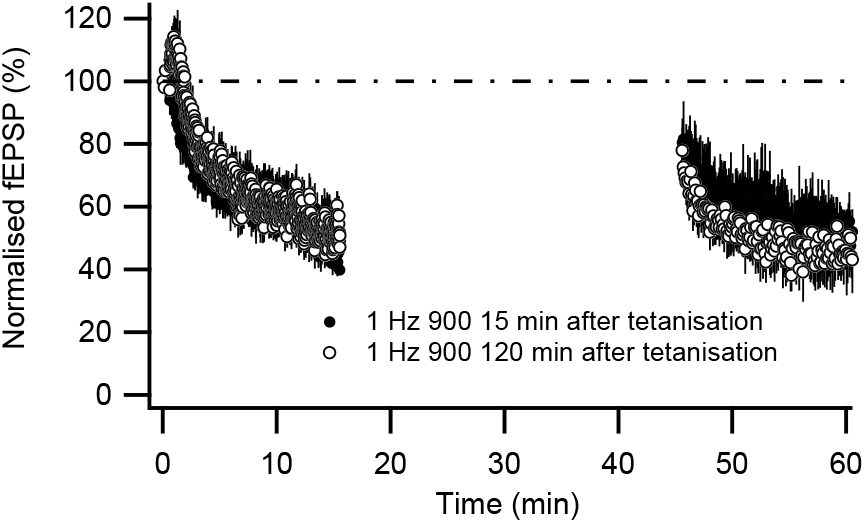
The effect of tetanization on 1 Hz-induced depression is not dependent on time after tetanization. 900 stimuli at 1Hz were given 15 min (n = 5 experiments, 1 experiment is lacking data after the 30 min stimulus interruption) or 120 min (n = 5 experiments, 1 experiment is lacking data after the 30 min stimulus interruption) after our standard tetanization protocol was applied (see text-Fig. 1 A).

## DISCUSSION

The present study shows that Hebbian activity does not stabilize AMPA signalling in developing CA3-CA1 synapses. Thus, even after strong repeated high frequency tetanization test pulse stimulation (1/5s) depresses the field EPSP far below its naive pre-tetanus level without a stable level of AMPA signalling being reached even after 3-4 hours of test pulse stimulation. This result contrasts with the stable potentiation observed in the mature hippocampus after 30-60 min of stimulation at similar rates of test pulse stimulation (e.g. 1/7.5 s, (Volianskis & Jensen, 2003)). Thus, the stabilizing action of Hebbian activity against test pulse induced depression noted in earlier reports (Abrahamsson et al., 2008; Xiao et al., 2004) is only temporary. We also found that the tetanization-induced transient potentiation above the naive AMPA signalling level did not decline if test pulse stimulation was suspended. High frequency tetanization thus leads to two seemingly disparate effects on the developing synapses; i) to a de-depression of any preceding test pulse induced depression back to the naive field EPSP level, and ii) to a potentiation exceeding the naive level. Both these effects are long-lasting if the synapses are presynaptically silent and are transient if the synapses are exposed to test pulse stimulation.

### Are there any AMPA stable CA3-CA1 synapses in the 2^nd^ postnatal week?

Since test pulse induced depression in the 2^nd^ postnatal week appeared to saturate at about 40% from the naive level it was suggested that the CA3-CA1 synapse population could be subdivided into AMPA labile and stable AMPA-mature synapses, the latter synapses stabilized by prior participation in Hebbian activity (Abrahamsson et al., 2008; Groc et al., 2006). We showed that more prolonged test pulse stimulation (900 stimuli at 0.05-0.2 Hz) resulted in depression to ∼ 60% from the naive level, and we now demonstrate that further 0.2 Hz stimulation (2700 stimuli in total) results in depression to ∼ 80% from the naive level. Thus, while this further depression might involve synapses less AMPA labile than those silenced by briefer test pulse stimulation, this large depression nevertheless suggests that a subdivision into AMPA stable and AMPA labile synapse in the developing hippocampus is no longer tenable. Furthermore, as shown presently, Hebbian activity does not create AMPA stable synapses at this developmental stage.

Application of the protein kinase A (PKA) activator forskolin was previously found to de-depress an on-going test pulse induced depression and to stabilize the field EPSP at the naive level (Abrahamsson et al, 2008). Such an action by forskolin agrees with the demonstration that tetanization-induced “LTP” in the 2^nd^ postnatal week relies on PKA activation (Yasuda et al., 2003) and that GluR2_long_-containing AMPARs at this developmental stage can be synaptically incorporated via NMDAR/PKA activation (Esteban et al., 2003; Kolleker et al., 2003; Qin et al., 2005; Zhu et al., 2000). Our results would thus suggest that any such NMDAR/PKA activation even by strong repeated high frequency tetanization of these synapses has no long-term stabilizing action on the AMPA signaling.

### Tetanization-induced potentiation above the naive level

While the present study shows that tetanization does not stabilize AMPA signaling it does indicate that the synapses become less labile. The evaluation of such a partial stabilizing effect of Hebbian activity is however hampered by our poor understanding of what underlies the potentiation exceeding the naive level and its lability compared to that of the pre-existing AMPA signaling. Such a potentiation has previously been observed in the mature hippocampus and thought to underlie the so called short-term potentiation (STP) (Volianskis & Jensen, 2003), and has been attributed to a presynaptic modification (Volianskis & Jensen, 2003). While we at present cannot exclude such a mechanism also in the developing hippocampus it has been observed, albeit in somewhat older animals (P14) than used here, that whereas LTP in GluR1 knock-out mice was not much less than in wild-type mice the STP was absent (Jensen et al., 2003). GluR1 subunits are abundantly present in CA1 pyramidal cells also in the 2^nd^ postnatal week (Li et al., 2003; Zhu et al., 2000) but require CaMKII, which is present only at a low level at this early time period (Kelly et al., 1987), for their stable synaptic insertion (Esteban et al., 2003; Hayashi et al., 2000). However, the lack of STP in GluR1 knock-out mice may suggest that NMDAR/PKA activity may insert GluR1-containing AMPARs into the synaptic membrane but only in a labile manner such that these receptors are removed by test pulse stimulation. Application of the PKA activator forskolin also results in a transient potentiation above the naive level (Abrahamsson et al., 2008). Both the de-depression to the naive level and the potentiation above that level may thus be explained by an NMDAR/PKA mediated insertion of AMPARs into the synaptic membrane, GluR2_long_–containing and GluR1-containing AMPARs, respectively.

### Comparison between depressions of naive vs tetanized synapses

If we assume that the potentiation above the naive level is as labile as indicated by its decay after a single tetanization event (<10-15 min) (Fig. 1A) (see also (Abrahamsson et al., 2008)), all subsequent depression would be only that of the pre-existing AMPA signaling and be evaluated from the naive level. Hebbian activity would thus appear to result in a substantial partial stabilization of both test pulse and 1 Hz-induced depression (Figs. 2 and 3). For example, after 2700 stimuli at 0.2 Hz the depression at tetanized synapses is 63% from the naive level compared to 82% at naive synapses, indicating that about twice as much of the pre-existing AMPA signaling remains after this number of stimuli at tetanized than at naive synapses. However, at the other extreme, we could assume that the potentiation above the naive level is as labile as the pre-existing AMPA signaling. This assumption has also the advantage that it compares the overall lability of AMPA signaling after a strong Hebbian induction vs that of naive synapses. The depression of the tetanized synapses should then be evaluated from the post-tetanus-naive level. Evaluated in this manner there was after 2700 stimuli no difference in lability between tetanized and naive synapses (Figs 2B and 3C). Furthermore, despite using this assumption that may overestimate the lability of the pre-existing AMPA signaling, the depression of tetanized synapses measured at earlier time points was less than the depression of naive synapses. It thus seems safe to conclude that tetanization does cause some stabilization of the pre-existing AMPA signalling, although to what extent this occurs is uncertain.

### Comparison between long-lasting depressions of naive vs tetanized synapses

We previously showed that 900 stimuli delivered at both 0.2 and 1 Hz resulted in a long-lasting depression of ∼ 40% from the naive level (Strandberg & Gustafsson, 2011). We presently found that additional 1800 stimuli at 0.2 Hz substantially increased this depression to 66% from the naive level suggesting that prolonged test pulse stimulation can result in long-lasting silence in about two-thirds of the CA3-CA1 synapses. After tetanization this long-lasting depression was found to be substantially reduced, at least as evaluated from the naive level (Fig. 2A). We believe however that this procedure leads to an underestimation of the long-lasting depression at tetanized synapses since the potentiation as observed after a single tetanization event is fully recuperated after 20 min of stimulus interruption after having been de-potentiated by 10 min of 0.2 Hz stimulation (Abrahamsson, Gustafsson and Hanse, unpublished observations). If we make the conservative assumption that after 2700 stimuli the de-potentiation of the potentiation (above the naive level) is reversed to much the same extent as the depression of the pre-existing AMPA signalling the long-lasting depression of tetanized synapses should instead be evaluated from the post-tetanus-naive level. Adopting this procedure, the long-lasting depression at tetanized synapses did not in relative terms differ from that at naive synapses neither using 0.2 Hz (Fig. 2B) nor using 1Hz stimulation (Fig. 3C), indicating that Hebbian activity does not affect the ability of 2700 stimuli to result in long-lasting depression.

Nonetheless, when the briefer stimulation of only 900 stimuli at 1 Hz was used the long-lasting depression was substantially smaller than that observed at naive synapses. Moreover, when measured at earlier time points the depression was smaller at tetanized than at naive synapses (Fig. 2B). Thus, Hebbian activity at least results in a temporary partial stabilization against depression. An inhibitory influence of a preceding LTP on LTD induction has previously been described in older animals as a time-delimited effect, preventing LTD induction within 60 min after LTP induction (Montgomery & Madison, 2002; Peineau et al., 2007). For example, using organotypic slice cultures Montgomery and Madison described protection from depression of recently unsilenced synapses lasting for about an hour. In the present study we found that the depression/long-lasting depression produced by 900 stimuli at 1 Hz was similar whether the stimulation was applied 15 min or 2 hours after tetanization. The presently observed stabilizing action of Hebbian activity thus remained if the synapses were not stimulated and it dissipated with prolonged stimulation.

### Prolonged 0.2 Hz stimulation is more effective than 1 Hz stimulation to produce long-lasting depression at naive synapses

While 2700 stimuli resulted on average in a larger depression when 1 Hz than 0.2 Hz was used, the 1 Hz stimulation resulted in a significantly larger reversal and in a smaller long-lasting depression. Moreover, after the stimulus interruption the reversed depression did not readily return to the pre-stimulus interruption level when stimulation was resumed. We previously showed that while NMDARs are not necessary for inducing depression during on-going stimulation they are both necessary and sufficient for inducing the long-lasting depression (Strandberg & Gustafsson, 2011). Since 1 Hz stimulation results in LTD of NMDAR-mediated synaptic transmission (Selig et al., 1995; Xiao et al., 1994) a possible explanation might be that the prolonged 1 Hz, but not 0.2 Hz, stimulation of developing synapses results in such an NMDA LTD of sufficient magnitude to impair induction of the long-lasting depression. The reduced depression observed when the stimulation was resumed after stimulus interruption is also similar to that observed when NMDARs are blocked specifically during this time period (Strandberg & Gustafsson, 2011), consistent with a 1 Hz-induced prolonged depression of NMDAR-mediated transmission.

### Functional considerations

The glutamate synapse appears to acquire its AMPARs early in an NMDAR independent manner, but the AMPARs may easily be lost upon synaptic activation. In the first postnatal weeks the main role for Hebbian activity may therefore be to stabilize AMPA signalling in the synapses that partake in such combined pre- and postsynaptic activity. The present result shows that in the 2^nd^ postnatal week CA3-CA1 synapses that participate in Hebbian activity are initially less easily depressed when exposed to low frequency activity. Nevertheless, even after extensive participation in Hebbian activity they do not in the long run appear significantly more stable than before. To maintain its AMPA signalling the synapse must then either be presynaptically silent, or continuously participate in Hebbian activity, synaptic activity outside this context leading to AMPA silencing and possible elimination. This would allow for a dynamic build-up and refinement of the synaptic circuitry during development, only allowing synapses that throughout this early developmental period are activated in proper relation to other synapses to survive.

